# Gut Microbiome Composition Reveals the Distinctiveness between the Bengali people and the Indigenous Ethnicities in Bangladesh

**DOI:** 10.1101/2023.02.15.528648

**Authors:** Ishtiaque Ahammad, Anisur Rahman, Zeshan Mahmud Chowdhury, Arittra Bhattacharjee, Gourab Dewan, Shiny Talukder, Keshob Chandra Das, Chaman Ara Keya, Mohammad Uzzal Hossain, Md. Salimullah

## Abstract

Human gut microbiome is influenced by ethnicity and other factors. In this study, we have explored the gut microbiome of Bengali population (n=13) and four Tibeto-Burman indigenous communities-Chakma (n=15), Marma (n=6), Khyang (n=10), and Tripura (n=11) using 16S rRNA amplicon sequencing. A total of 19 characterized phyla were identified in 5 cohorts, with *Firmicutes* and *Bacteroidetes* being the most prevalent. At the genus level, the abundance of *Prevotella* was relatively similar across all ethnicities. However, the Chakma population demonstrated higher *Bacteroides* abundance. Chakma people were more distinct than other ethnicities and exhibited a higher quantity of differentially abundant microbial features. The Bengali population had relatively low bacterial richness and *‘Firmicutes* to *Bacteroidetes* ratio’ than others with lower qualitative microbial diversity. A phylosymbiotic link between Bangladeshi indigenous people and certain ethnic groups in India have also been discovered. A comparative analysis between all Bangladeshi samples (n=55) and several tropical and subtropical countries (n=132) such as Australia, Egypt, Indonesia, Malaysia, Mexico, Thailand, and Vietnam revealed that the gut microbiota profile of Bangladeshi people is remarkably distinct from others. The insights from this study will aid further epidemiological and translational research.

## Introduction

The human body harbors trillions of microorganisms that outnumber human cells by a large margin and contains genes unique to human genomes^1^. The gastrointestinal tract possesses a wide range of microbes that assist physiological functions such as maintenance of the gut barrier, immunomodulation, and nutrient metabolism ^2,3^. Several studies have shown that gut bacteria are essential for maintaining human health ^4–6^. Based on culture-dependent and independent techniques, the gut microbiome of any human individual is estimated to contain 150 to 400 species. The majority of these species belong to four phyla: *Bacteroidetes, Firmicutes, Actinobacteria*, and *Proteobacteria*^5^ Although gut microbiomes vary at the individual level, several recognizable patterns have been observed in the population of western and non-western countries ^7–10^. Imbalance in the gut microbiota has been linked to a variety of diseases such as inflammatory bowel disease, obesity, allergies, and psychological disorders ^11,12^.

Earlier studies have suggested that factors such as unfavorable diet, environment, age, and lifestyle can promote the gut microbiome imbalance^8,13,14^. However, the extent to which each of these factors regulates the microbiome is disputed. Previous studies showed that the relative abundance of common phylum *Bacteroides* is higher in Western gut microbiomes, whereas *Firmicutes* and *Proteobacteria* are more prevalent in non-Western gut microbiomes^15,16^. Moreover, the process of industrialization exerts a significant impact on the composition of the gut microflora. For instance, people in Africa and South America leading a traditional lifestyle share a higher diversity of microbes compared to industrialized countries in Europe and North America^7^. Hence, different cultures around the world with variations in regular diet, sanitation, and medical practices impose a noticeable impact on the gut microbiota. It is reported that human communities that live in close vicinity yet practice different types of food habits have slight variation in alpha diversity across lifestyles. However, macroecological biodiversity, climate, and a variety of other characteristics must also be considered^17–19^. Therefore, examining communities that live in similar geographic regions and have undergone recent changes in culture, lifestyle, and diet would help researchers better understand the causal factors that affect the human gut microbial composition. These types of observational studies can be carried out using metagenomics-based high-throughput sequencing.

High-throughput metagenomic sequencing data involves two processes, such as the Amplicon-based method that uses 16S ribosomal RNA for bacteria, the internal transcribed spacer (ITS), and the 18S region for fungi and eukaryotes, respectively, or whole metagenomic shotgun sequencing. Amplicon-based 16S rRNA sequencing is a frequently used method for making community-wide taxonomic designations depending on the variable sections (V1-V9) of the bacterial *16S rRNA* gene^20^. It has been used to analyze microbial diversity in a variety of environmental samples, including soil and human gut^21,22^. On the other hand, shotgun metagenomic analysis involves both sequence-based and functional screening, revealing microbial diversity, genomes, and functional gene products without stating the origin of the genetic material^23^. Additionally, getting proper coverage and depth for species detection makes this technique more expensive than others^24,25^.

The Indigenous population in Bangladesh consists of 54 indigenous groups, accounting for 1.8 percent of the overall population of the country while the rest are Bengali^26^. Bengali people inhabit all over Bangladesh and depend on a rice-based high-carbohydrate diet with less nutritional diversity^27^. On the other hand, the Chakma, Marma, Khyang, and Tripura, the major indigenous groups in Bangladesh, majorly reside in the tropical Chittagong hill tracts (Rangamati, Bandarban, and Khagrachari districts) of southeastern Bangladesh. They share astonishingly high genetic homogeneity among themselves, as well as substantial similarities with the Tibeto-Burman population in Northeast India. They also have a large proportion of genetic descent from the mainland of India^28^. Though these indigenous communities are increasingly adopting the modern education and lifestyle of the majority Bangladeshi population, yet each group has largely preserved its own culture and food habits. Most of them take rice as their staple food, which is often supplemented by fish, meat, eggs, maize, millet, and various vegetables like yams, melons, pumpkins, and cucumbers^29^. Besides, each tribe has its traditional dishes, e.g., for example, the Chakma people consume bamboo shoots, while the Marma love to eat “Nappi,” a dried fish paste. They also smoke indigenous cigarettes and drink home-brewed fermented beverages. As each of these cohorts has a differing ethnicity and cultural tradition, it is more likely that there will be perceptible differences in microbial compositions in their gut.

In this study, the impact of ethnicity on the gut microbiota was investigated. 16s rRNA sequencing was used to find distinguishable patterns in the gut microbial community between Bengali and four indigenous cohorts as well as any existing differences between the indigenous tribes themselves. Besides, several publicly available foreign datasets were retrieved from the NCBI for comparative analysis with Bangladeshi samples sequenced in this study.

## Material and Methods

### Sample Collection

Prior to sample collection, written informed consent was obtained from all participants. Ethical approval and permits were obtained from the National Institute of Biotechnology Ethical Review Committee, Bangladesh (NIBERC2022-01). **Supplementary Table 1** contains detailed sample information on age, gender, and cohort.

All of the indigenous volunteers in this study reside in the Rangamati district (coordinates: 22°37’60N 92°12’0E; Altitude: 15 meters above sea level) of the Chittagong Hill Tracts region. Bengali samples were collected from Dhaka Division (23.9536° N, 90.1495° E; flat lands). Based on ethnicity, the participants were divided into five groups (Bengali, Chakma, Marma, Khyang, and Tripura). Fecal samples of the participants were collected using Sterile stool collection tubes. The samples were immediately placed in iceboxes and transported to the laboratory where they have been stored at −80 °C temperature. Fecal DNA extraction was executed using the PureLink™ Microbiome DNA Purification Kit (Catalog number: A29790). Specialized beads were used along with a combination of heat, chemical, and mechanical disruption to lyse the microorganisms efficiently. Precipitation with a proprietary cleaning buffer removed inhibitors. After that, the samples were placed in spin columns, and the DNA attached to the column was washed once before elution. DNA concentration and purity were estimated by Thermo Scientific NanoDrop 2000/2000c and then stored at −20°C.

### Amplicon Generation and Library Preparation for Sequencing

Amplicons were generated using the 341F (5’-CCTAYGGGRBGCASCAG-3’) and 806R (5’-GGACTACNNGGGTATCTAAT-3’) primers that targeted the V3-V4 region of the bacterial *16S rRNA* gene. All PCR reactions were performed with 15 μL of Phusion^®^ High-Fidelity PCR Master Mix (New England Biolabs), 0.2 μM of forward and reverse primers, and around 10 ng template DNA. For thermal cycling, initial denaturation at 98°C for 1 minute was followed by 30 cycles of denaturation at 98°C for 10 seconds, annealing at 50°C for 30 seconds, and elongation at 72°C for 30 seconds. Final elongation was carried out for 5 minutes at 72°C. TruSeq^®^ DNA PCR-Free Sample Preparation Kit (Illumina, USA) was used as per the manufacturer’s protocol for sequencing library preparation, and index codes were appended. The quality of the library was evaluated by employing the Qubit@2.0 Fluorometer (Thermo Scientific) and the Agilent Bioanalyzer 2100 system. Finally, the library was sequenced on an Illumina NovaSeq platform, resulting in 250 bp paired-end reads.

### Data Pre-processing and Quality Control

The data pre-processing and most of the analysis were performed through different tools (q2) of the QIIME2 (version 2021.4.0) command-line interface^30^. In addition, essential graphs and plots were further made using various R packages from the tidyverse package collection, such as ggplot2, tidyr, dplyr, and so on^31^. Raw data from the sequencing platform was de-multiplexed into individual samples based on their unique barcodes and PCR primers were removed by the q2-cutadapt tool^32^. For quality control of reads, the DADA2 R library was implemented as a QIIME2 tool that filters noisy sequences, performs error correction in marginal sequences, removes chimeras and singletons, joins denoised paired-end reads, and also dereplicates the filtered reads^33^. The features produced by dada2 are denoted as amplicon sequence variants (ASVs).

### Taxonomic Assignment

A naive Bayes classifier was trained by using Greengenes latest 13_8 reference dataset based on 99% sequence similarity^34,35^. Here, the q2-feature-classifier tool from the Qiime2 command-line interface was deployed to train the classifier^36^. If study samples are sequenced by targeting a specific variable region of the *16S-rRNA* gene, a naive Bayes classifier should also be trained on the reference sequences extracted from the same region. As it has been demonstrated that such targeted training of classifiers helps to attain higher taxonomic resolution^37^. In the current study, the taxonomic classifier was trained on the reference sequences from the V3-V4 region of the 16S-rRNA gene as the samples of this study were sequenced by targeting this region. This classifier was then deployed for pre-fitted sklearn-based taxonomy assignment of ASVs (also denoted as features) previously generated by dada2. To depict the taxonomic composition, a box plot and a krona plot have been generated by the q2-taxa tool and KronaTools (version 2.7.1) respectively^38^. The features present within the 0.9 fractions of the samples (in 49 samples) were considered to be core features. These features were identified by the core-features command of the q2-feature-table tool. The frequency heatmap of these core features was plotted by the heatmap command of the q2-feature-table tool of QIIME2^39^. In this instance, the “Euclidean” metric and “Average” method were attributed to the log10 transformed normalized data^40^.

### Diversity Analysis

Several diversity metrics in QIIME2 require a rooted phylogenetic tree generated from the ASVs of the sampled data. A reference-based fragment insertion method was applied to construct the rooted tree for this purpose. Greengenes 13_8 data from the SEPP-Refs project (https://github.com/smirarab/sepp-refs/) was used as a reference database in the q2-fragment-insertion tool^41,42^. The sequencing depth of the samples needed to be enough so that the richness of the samples could be fully observed. This phenomenon was checked with the alpha rarefaction curve generated by the q2-diversity tool. The microbiome within and between samples was calculated by the core-metric-phylogenetic method of the q2-diversity tool. This method computes several alpha (Observed features, Shanon diversity, Faith’s phylogenetic diversity, Pielou’s evenness) and beta diversity (Jaccard distance, Bray-Curtis distance, unweighted UniFrac distance, and weighted UniFrac distance) metrics altogether^43^. Based on each beta diversity metric, this command also performed principal coordinates analysis (PCoA)^44^. EMPeror visualization tool, as an integrated tool in QIIME2, was utilized to create the PCoA plots for every beta diversity metric^45^.

The Kruskal–Wallis H test was performed by the alpha-group-significance command of the q2-diversity tool to visually and statistically assess differences in alpha diversity between distinct groups of samples^46,47^. In many cases, several interacting factors together can impact the alpha diversity. The two-way ANOVA test was conducted (by the q2-longitudinal tool) to check whether the interaction of the Cohort category along with other factors (Sex and Age_group) had a combined effect on alpha diversity^48^. This test was also followed by a paired t-test for each alpha diversity metric. The beta-group-significance command of the q2-diversity tool was then employed to perform the PERMANOVA test to investigate if samples from the same ethnicity are more similar to each other than samples from different tribes^49^.

### Differential Abundance Test

To classify the features that were differentially abundant across various sample groups the analysis of the composition of microbiomes (ANCOM) method was applied by the q2-composition tool^50^. This statistical framework was deployed at the species level of the features and pseudo count was added to the features with zero frequencies. The minimum sample size for each feature was set to 27 (half of the total samples) because ANCOM fails to manage false discovery rates at sample sizes less than 10 as well as to remove very lowly abundant features^51^. For linear discriminant analysis (LDA) by the LEfSe, the feature table was collapsed at the genus level and the minimum sample size was chosen at 27^52^. This tool first performs the non-parametric Kruskal-Wallis (KW) sum-rank test to identify the features which had significant differential abundance across different metadata categories. Finally, LEfSe applies LDA to compute the effect size of each differentially abundant feature and plot the LDA score in the log10 scale.

### Functional Analysis

BURRITO, an interactive visualization web server (http://elbo-spice.cs.tau.ac.il/shiny/burrito/), was utilized to explore the taxa-function relationship within the samples of the study^53^. To acquire gene contents and functional annotations, this tool adopts the PICRUSt and KEGG Orthology databases, respectively^54,55^. At first, features with a sample size of less than 27 were filtered from the original feature table to remove very low abundant taxon. The q2-vsearch tool was employed for closed-reference clustering of retrieved features at 97% identity based on greengenes 97% OTU IDs as reference^56,57^. Thus the acquired OTU table was then converted to the appropriate table format for BURRITO input.

### Phylosymbiosis Analysis

The 16S metagenomic data of other rural and urban populations around the world were obtained from the MG-RAST server to compare them with the gut microbiota of the people of Bangladeshi origin^58^. These foreign datasets included 75 samples from 15 tribal populations from 4 geographic regions (Telangana, Sikkim, Assam, Manipur) of India, 22 samples from Mongolia, 8 samples from Amerindian people of Venezuela, 13 Malawian samples, and 14 American samples^8,13,59^. Relevant MG-RAST IDs for each sample are documented in **Supplementary Table 2**. After importing all data separately as QIIME2 artifacts, quality control, and filtration was performed by q2-dada2, q2-deblur, and q2-quality-filter tools^33,60,61^. The choice of different tools and commands for denoising and ASVs generation was based on sequencing types and amplicon lengths of each study. As the data from different sources had different sequencing depths, all samples were rarefied at 300000 depth by sampling with replacement. Then all feature tables and representative ASVs were united with the merge and merge-seqs command of the q2-feature-table tool of QIIME2. Subsequently, a phylogenetic rooted tree was generated by the sepp fragment insertion approach. This rooted phylogeny was used to create a UPGMA (Unweighted Pair Group Method with Arithmetic Mean) tree based on the Unweighted UniFrac metric by beta-rarefaction command of the q2-diversity tool^62,63^. The phylogenetic tree was drawn and annotated using the iTOL online visualization tool^64^. Additionally, principal coordinate analysis (PCoA) was also conducted and plotted based on both Unweighted and Weighted UniFrac metrics to visualize the beta diversity amidst the collected samples^44^.

### Comparative Analysis with Tropical and Sub-tropical Countries

To compare the gut microbiome composition of Bangladeshi samples in our study with tropical and sub-tropical countries, publicly available 16s rRNA amplicon sequence data was retrieved from the NCBI Sequence Read Archive (NCBI-SRA). Only healthy / control samples were taken from the selected datasets. Countries included in the comparative analysis were Australia, Egypt, Indonesia, Malaysia, Mexico, Thailand, and Vietnam. The number of samples taken and NCBI project IDs for these datasets have been presented in **Table 1**. These foreign samples (n=132) were then compared with all Bangladeshi samples (n=55) using the q2-diversity tool. Alpha diversity of all Bangladeshi and the tropical and sub-tropical samples was measured based on Shannon’s diversity and Faith’s phylogenetic diversity index. For comparison of beta diversity, community dissimilarity was measured qualitatively and quantitatively using Bray-Curtis distance and Jaccard distance methods respectively.

**Table 1:** Publicly available 16s gut microbiome datasets of tropical and sub-tropical countries downloaded for comparison with the Bangladeshi samples.

| Country | NCBI_BioProject_ID | Reference | No_of_samples |
| --- | --- | --- | --- |
| Australia | PRJNA393083 | <sup>107</sup> | 30 |
| Egypt | PRJNA629382 | <sup>108</sup> | 7 |
| Indonesia | PRJDB9293 | <sup>109</sup> | 23 |
| Malaysia | PRJNA631204 | <sup>110</sup> | 30 |
| Mexico | PRJNA719138 | <sup>111</sup> | 23 |
| Thailand | PRJDB5860 | <sup>112</sup> | 30 |
| Vietnam | PRJNA668251 | <sup>113</sup> | 7 |

## Results

### Data Overview

A total of 55 individuals were sampled, of which 13 were Bengalis, 15 Chakmas, 10 Khyang, 6 Marma, and 11 Tripurans. After extracting DNA from fecal samples, the V3-V4 segment of the *16S rRNA* gene was amplified and sequenced. Demultiplexed barcodes and PCR primer removed raw data have been deposited in the NCBI database (Project ID). The sequence count per sample is listed in **Supplementary Table 3**. After conducting denoising, merging, and chimera-removal, q2-DADA2 discovered 6,809 features (ASVs) with 3,817,015 total frequencies in 55 samples. Overall denoising stats, feature table summaries, and representative sequences for each ASVs are provided in **Supplementary Table 4, 5, and, Supplementary File S1**.

### Bengali Population Possessed Lower Firmicutes to Bacteroidetes (F/B) Ratio Compared to Indigenous Groups

Among the 19 classified phyla, *Firmicutes* and *Bacteroidetes* were the most prevalent (48% and 34% of total) in the gut microbiota of all Bangladeshi populations. Other phyla with higher abundance were Proteobacteria (14%), Actinobacteria (3%), Tenericutes (0.5%), etc (**Supplementary Fig. 1**). Interestingly, 100% of taxa from the *Bacteroidetes* phylum belonged to the *Bacteroidia* class. On the other hand, considering *Firmicutes* phylum, 87% of features are *Clostridia* class, 11% *Bacilli*, and 2% are *Erysipelotrichi* (**Supplementary Fig. 3**). In **Supplementary File S2**, the taxonomic classification of each feature id is documented, along with the confidence value.

The normalized abundance of the top ten genera across different cohorts is represented in cpm (counts per million) (Fig. 1a). The *Prevotella* genus abundance was relatively similar in all cohorts. Nevertheless, the prevalence of the *Bacteroides* genus was drastically higher in the Chakma population and very low in Marma and Tripura samples. Moreover, Chakma tribal group also contained *Faecalibacterium*, *Roseburia*, and uncharacterized genera from the *Ruminococcaceae* family and *Bacteroidales* order in a relatively higher amount. On the other hand, *Streptococcus* and *Megasphaera* genera were extremely abundant in the Bengali and Tripura populations and the Marma cohort contained these genera in relatively lower quantities. The Khyang group contained the genus *Dialister* and uncharacterized genera from *Enterobacteriaceae* and *Ruminococcaceae* families. About 0.05% ASVs were classified as archaea of the *Methanobacteriaceae* family, of which only 0.1% were from the *Methanosphaera* genus and the rest of the features belonging to the Methanobrevibacter genus (**Fig. 1b**).

**Fig. 01:**
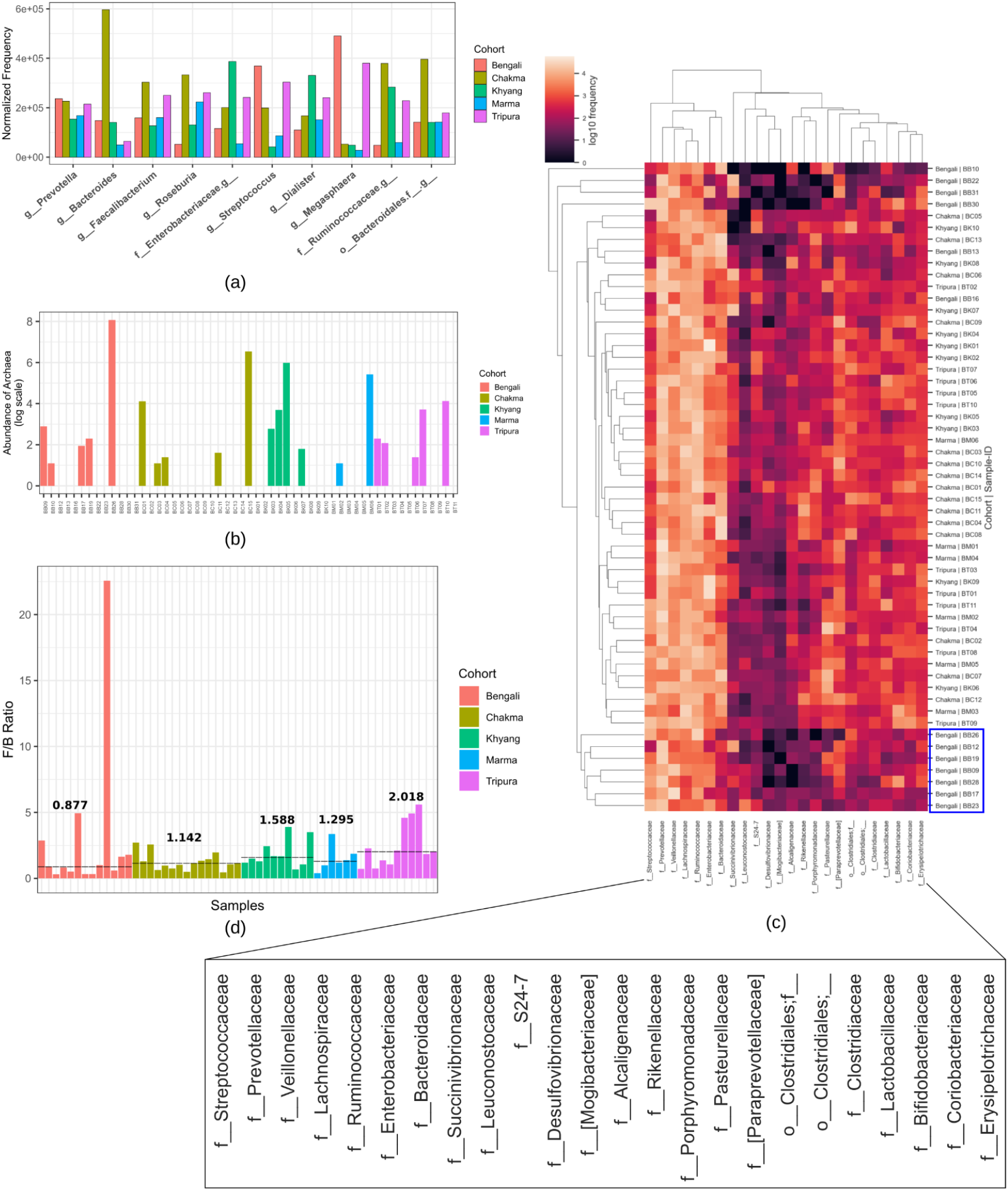
The abundance of the top prevailing bacterial genera, archaeal features, core features, and Firmicutes to Bacteroidetes ratio in various Bangladeshi cohorts. (a) The bar plot represents the normalized frequency of the top genera among different cohorts. The almost similar prevalence of the Prevotella genus in Bangladeshi cohorts reflects the country’s largely vegetarian diet. (b) Several archaeal species appeared to be present in different samples from each cohort. (c) The frequency of core features resulted in two primary clusters, with somewhat varied abundances in some Bengali samples. (d) The median ratio of Firmicutes to Bacteroidetes (F/B) implies that indigenous groups have a greater propensity for obesity.

Twenty-four families of features were found as core features and the frequency heatmap of these features revealed that there were two main clusters, one with higher frequency and another with lower frequency (**Fig. 1c**). The *Streptococcaceae, Prevotellaceae, Veillonellaceae, Lachnospiraceae, Ruminococcaceae, Enterobacteriaceae*, and *Bacteroidaceae* families were in the higher frequency cluster. Most of the samples from the Bengali population clustered together and contained these higher frequency microbial families in relatively lower amounts. The median value of the Firmicutes to Bacteroidetes (F/B) ratio was highest (2.018) in the case of Tripura samples, while the Bengali population had the lowest (0.877) median ratio **(Fig. 1d)**. All other cohorts (Chakma, Khyang, Tripura) had a median F/B ratio of more than one.

### The Qualitative Bacterial Diversity of the Bengali population was Lower than Indigenous Groups

The core-metric-phylogenetic method requires even sampling depth for calculation. Based on information from the frequency of features per sample and alpha-rarefaction curve 49561 was used as sampling depth, and 67.52% of features were retained at this depth. Three Bengali samples were excluded in this case due to not having enough sampling depth.

In the case of two alpha diversity metrics (Observed features and Shannon entropy), a significant difference was found within the samples (**Fig. 2**). Moreover, the Bengali samples had higher variation per individual in terms of bacterial richness, evenness, and phylogenetic relationships. Individuals within all indigenous cohorts exhibited very consistent phylogenetic diversity in microbial communities. In the case of the pairwise t-test in two-way ANOVA, the Bengali, and Chakma groups had a significant difference in both qualitative (Observed features) and quantitative (Shannon entropy) bacterial richness (BH FDR p-value: 0.036 and 0.015). Detailed results of other groups can be found in **Supplementary File S3**.

**Fig. 02:**
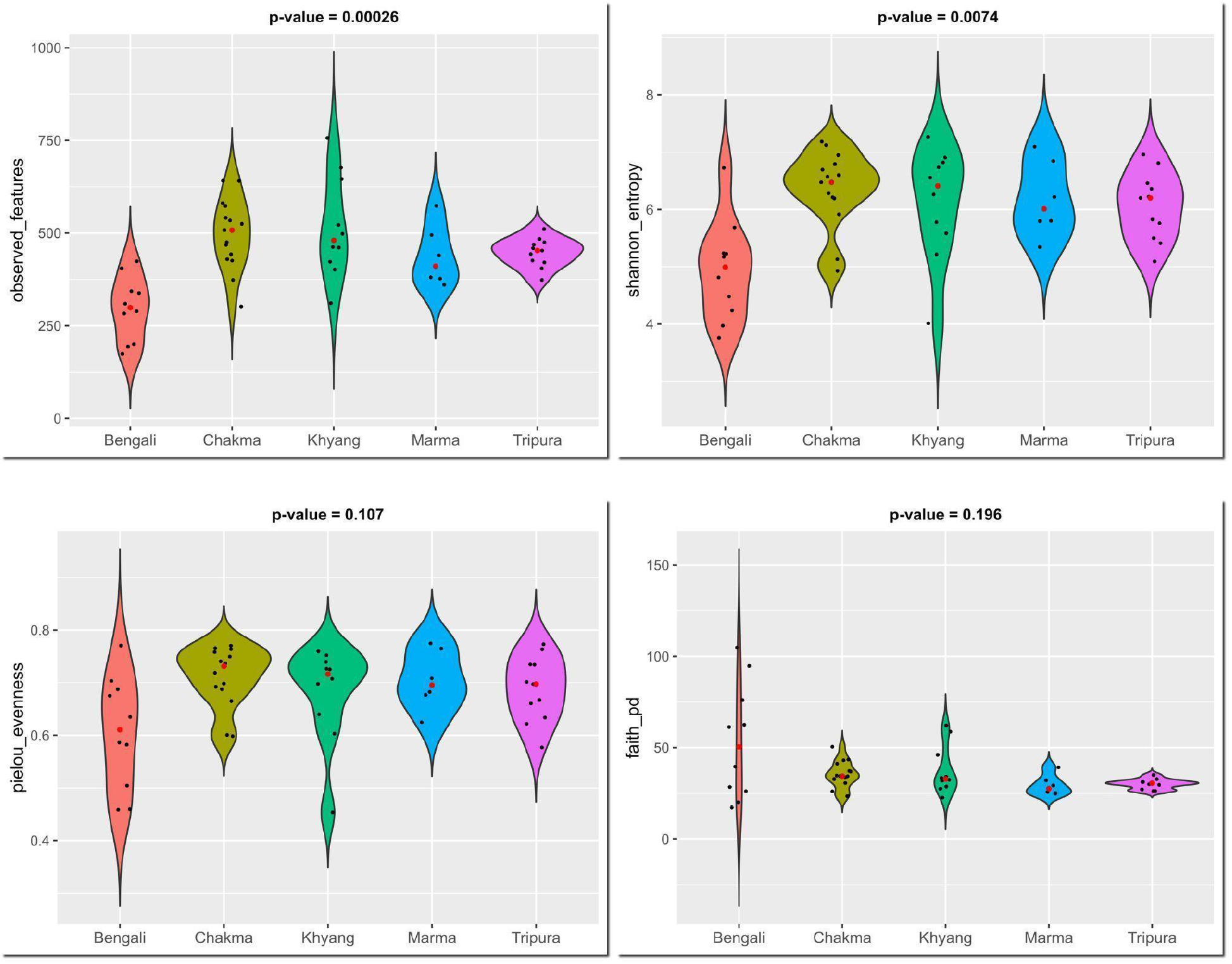
Distribution of alpha diversity among various Bangladeshi cohorts. For the Bengali, Chakma, Khyang, Marma, and Tripura populations there were statistically significant differences in alpha diversity values regarding qualitative and quantitative community richness (observed features and shannon entropy). Bengali samples exhibited a wide range of values across all alpha diversity metrics compared to other samples. The red dot inside each violin plot represents the median value for the corresponding cohort.

The PCoA plot revealed that the Bengali cohort was distinct from others in terms of qualitative metrics (Jaccard distance and Unweighted UniFrac distance) of bacterial community dissimilarity (**Fig. 3a and 3b**). **Fig. 3c** denotes the beta diversity difference between Cohorts based on the PERMANOVA test. The cohorts had significant differences in beta diversity in the case of Bray-Curtis distance (p-value 0.001) and Jaccard distance (p-value 0.001), Unweighted UniFrac distance (p-value 0.001), and Weighted UniFrace distance (p-value 0.026) metrics. But pairwise PERMANOVA statistics revealed that the Bengali samples had statistically significant beta diversity differences from the samples of other cohorts. Because, for all beta diversity metrics except Weighted UniFrace distance, Bengali vs Chakma, Bengali vs Marma vs Tripura, and Bengali vs Khyang pairwise test had FDR adjusted p-value (q-value) below significance level (**Fig. 3c**). Additionally, the Chakma cohort also had a significant difference in the quantitative measure of microbial community dissimilarity (Bray-Curtis distance).

**Fig. 03:**
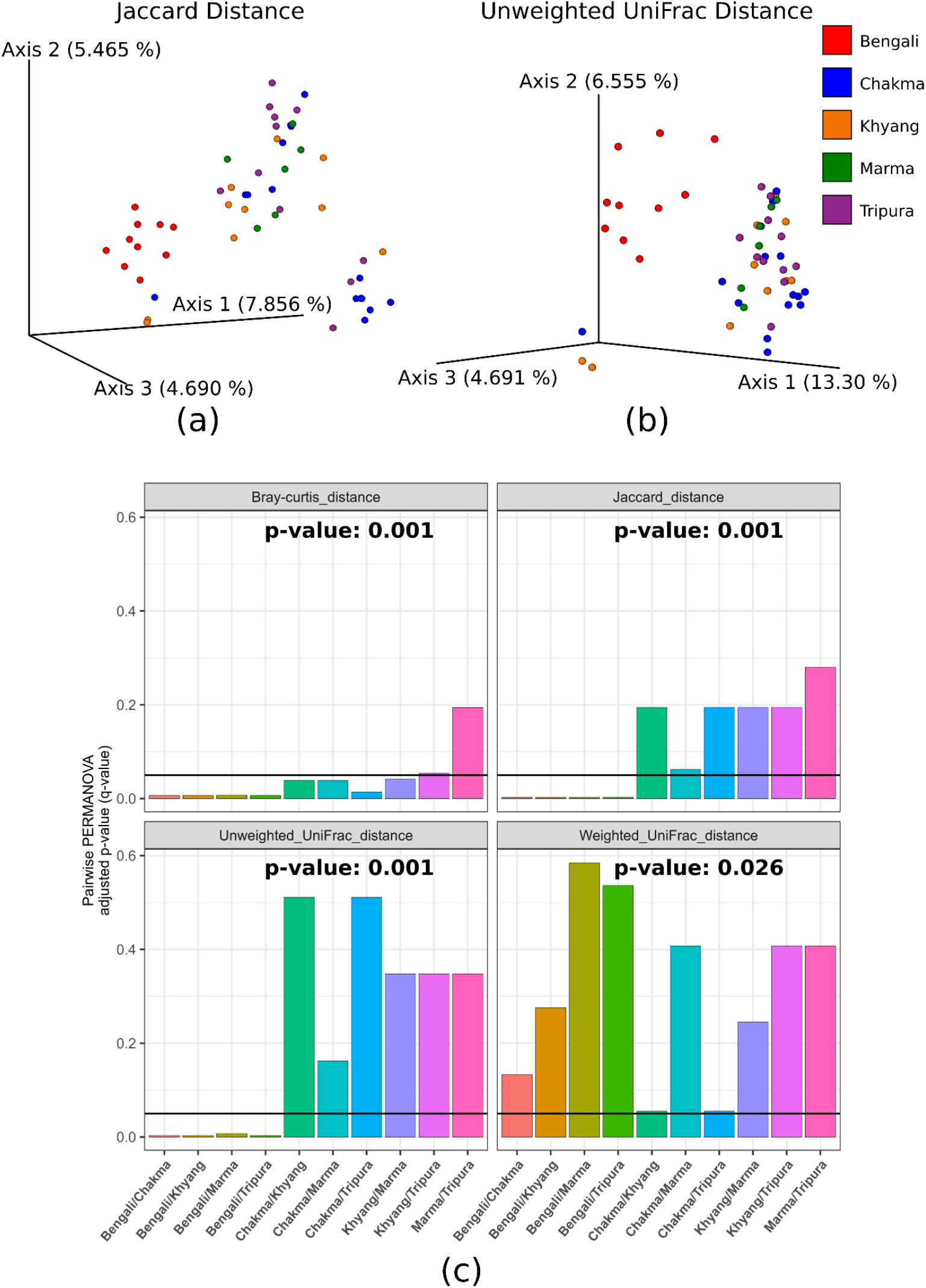
Principal coordinate analysis (PCoA) and the PERMANOVA test on the basis of beta diversity metrics. (a and b) In the case of qualitative measurements of beta diversity distance (Jaccard distance and Unweighted UniFrac distance), Bengali samples clustered together in the PCoA plot. (c) When the PERMANOVA statistics test was used to examine the beta diversity distance, it was shown that there were statistically significant differences between the cohorts. The black horizontal line indicates the statistical significance level of the p-value for pairwise PERMANOVA tests.

### The Chakma people harbored a richer bacterial community with differentially abundant species

In the ANCOM test for finding differentially abundant taxa between various sample groups, Chakma vs Non-Chakma and Bengali vs Non-Bengali comparison had differential taxa. The *Bacteroides caccae* bacteria had a higher abundance in Non-Bangali samples, and in contrast, an unclassified bacterial feature was highly abundant in Bengali samples **Fig. 4a**. In the case of the Chakma category, the taxa whose ANCOM statistics value (W value) was 8 or more, were plotted in **Fig. 4b**. Details ANCOM test results of this category are listed in **Supplementary File S4**. The *Prevotella stercorea* and an *Odoribacter* sp. were in relatively higher abundance in Chakma samples. *Bacteroides caccae, Anarostipes sp., Alistipes onderdonkii, Parabacteroides sp., Bacteroides uniformis, Bacteroides ovatus*, etc bacterial species had differentially lower abundance in the Chakma cohort (**Fig. 4b**). Linear discriminant analysis (LDA) by lefse up to the genus level yielded a large number of microbial features for each cohort (**Fig. 4c**). Every cohort seemed to be abundant with a wide variety of bacterial taxa (phylum to genus).

**Fig. 04:**
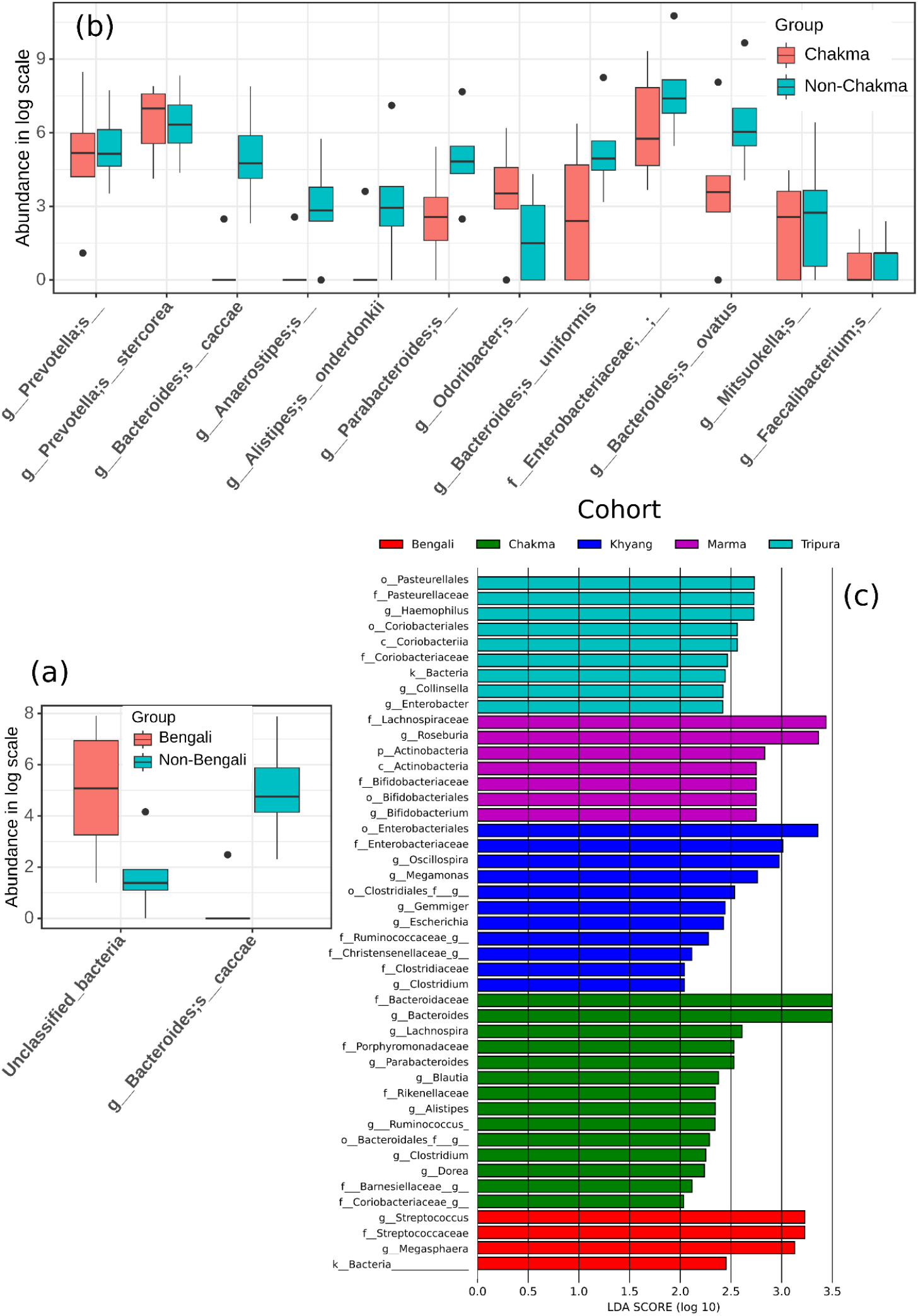
Differentially abundant bacterial characteristics as determined by ANCOM and LDA tests. (a) According to the ANCOM test, Bengali samples had a lower abundance of Bacteroides caccae bacteria than samples from other ethnic groups. (b) Comparing Chakma to other cohorts, the same statistical test produced a wide variety of bacterial species. (c) Finally, linear discriminant analysis (LDA) by LEfSe revealed a different number of differentially abundant bacterial taxa in each Bangladeshi cohort.

### Bengali Gut Microbiota Contributed to Markedly Diverse Biological Pathways

Relationships between microbiome composition and its effect on different biological functions were also explored in this study. Among all metadata categories, only the Bengali, Chakma, and Marma groups showed differential enrichment of function based on their gut microbiome composition **Fig. 5**. All differentially abundant functions along with related contributing taxa and BH (Benjamini and Hochberg) FDR adjusted p values are tabulated in the **Supplementary File S5**. A large number of pathways were differentially abundant in Bengali samples. Among them, the DNA repair system, Pyrimidine and purine metabolism, Lipopolysaccharide biosynthesis, peptidases, ribosome, etc functions had higher enrichment in Bengali samples. On the other hand, cell growth, transcription factors, transporters, bacterial motility protein, flagellar assembly, bacterial chemotaxis, etc functions had lower abundance among the Bengali population (**Fig. 5a**). The genus *Bacteroides, Blautia, Collinsella, Coprococcus, Dorea, Parabacteroides, SMB53, and Slackia* were the top significantly contributing taxa in the enrichment of these functions. Chakma samples seemed to be enriched with histidine metabolism, arginine biosynthesis, and transcription machinery (**Fig. 5b**). But, in the case of the Marma samples histidine metabolism, arginine biosynthesis functions were in relatively lower abundance than other Non-Marma samples, and transcription machinery function was in higher frequency (**Fig. 5c**). For both categories, the genus *Bacteroides, Blautia, and Parabacteroides* were the most contributing taxa for function abundance. Detailed statistics for significant pathways and taxa contributions for each pathway are documented in **Supplementary File S5**.

**Fig. 05:**
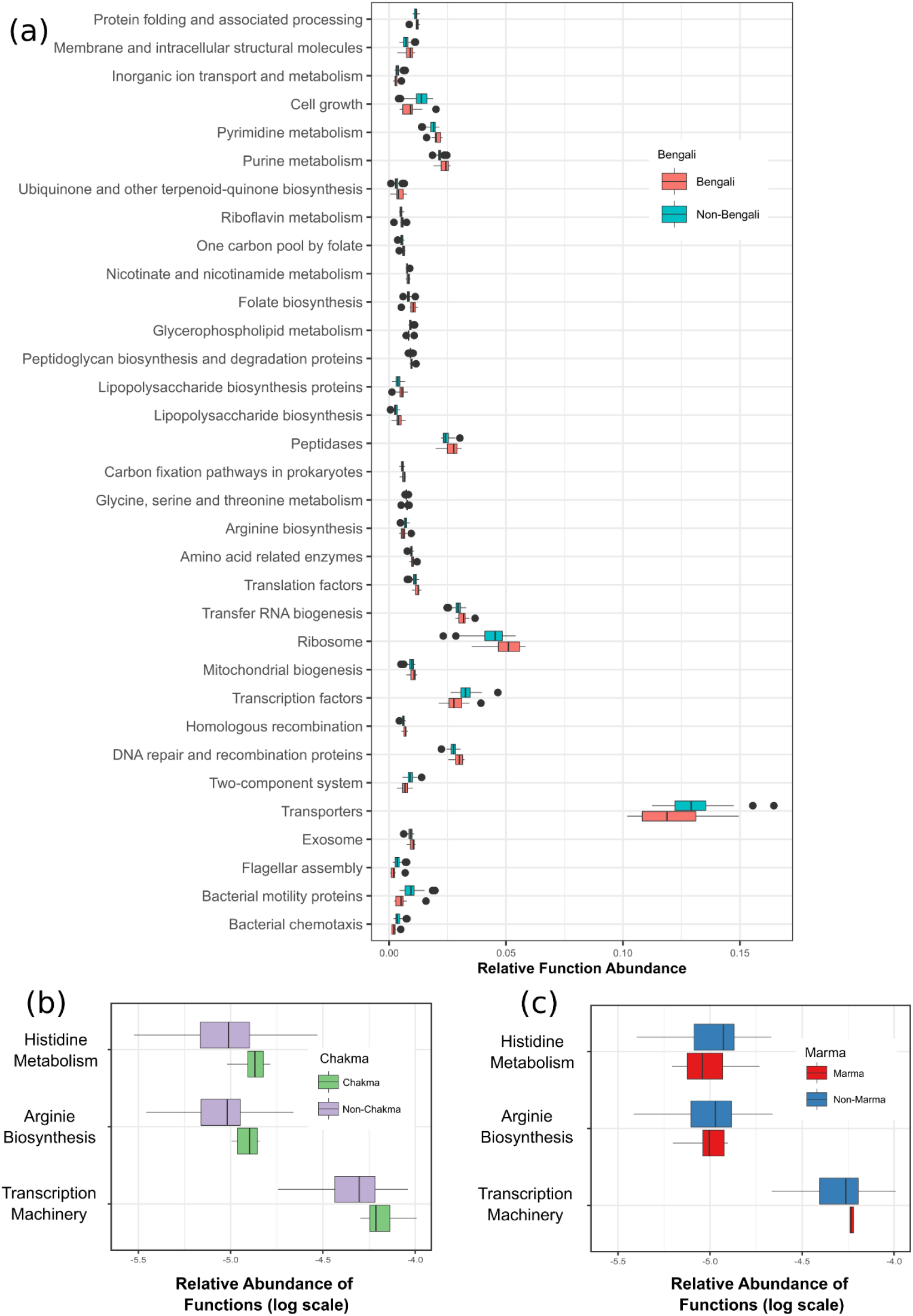
Pathway enrichment by bacterial communities among Bengali, Chakma, and Marma cohorts. (a) In comparison to other populations, the Bengali group appeared to have a broad range of differentially enriched pathways. Histidine metabolism, arginine biosynthesis, and transcription machinery, on the other hand, were differentially prevalent pathways in the Chakma (b) and Marma (c) cohorts.

### Bangladeshi Indigenous Tribes Exhibited a Similar Gut Microbiota Profile with North-east Indian Ethnic Groups

In the PCoA plot for the Unweighted and Weighted UniFrac beta diversity metric, Mongolian samples were clustered separately from other samples (**Fig. 6a**). The Bangladeshi samples clustered in between Indian and other-country (Malawian, American, Amerindian) samples in case of Unweighted UniFrac distance. But, when the phylogenetic quantitative measurements were added to the Weighted UniFrac distance, all samples were mixed except the Mongolian data (**Fig. 6b**). For beta rarefaction, the sampling depth was set to 300000. In the UPGMA tree, almost all Bangladeshi indigenous samples clustered in a separate branch but shared a common ancestral node with the Indian clades (**Fig. 6c**). Most of the Bengali samples clustered separately from Bangladeshi tribal samples. Some samples seemed to cluster on the distal branches, which can be considered outliers or affected by any internal factor.

**Fig. 06:**
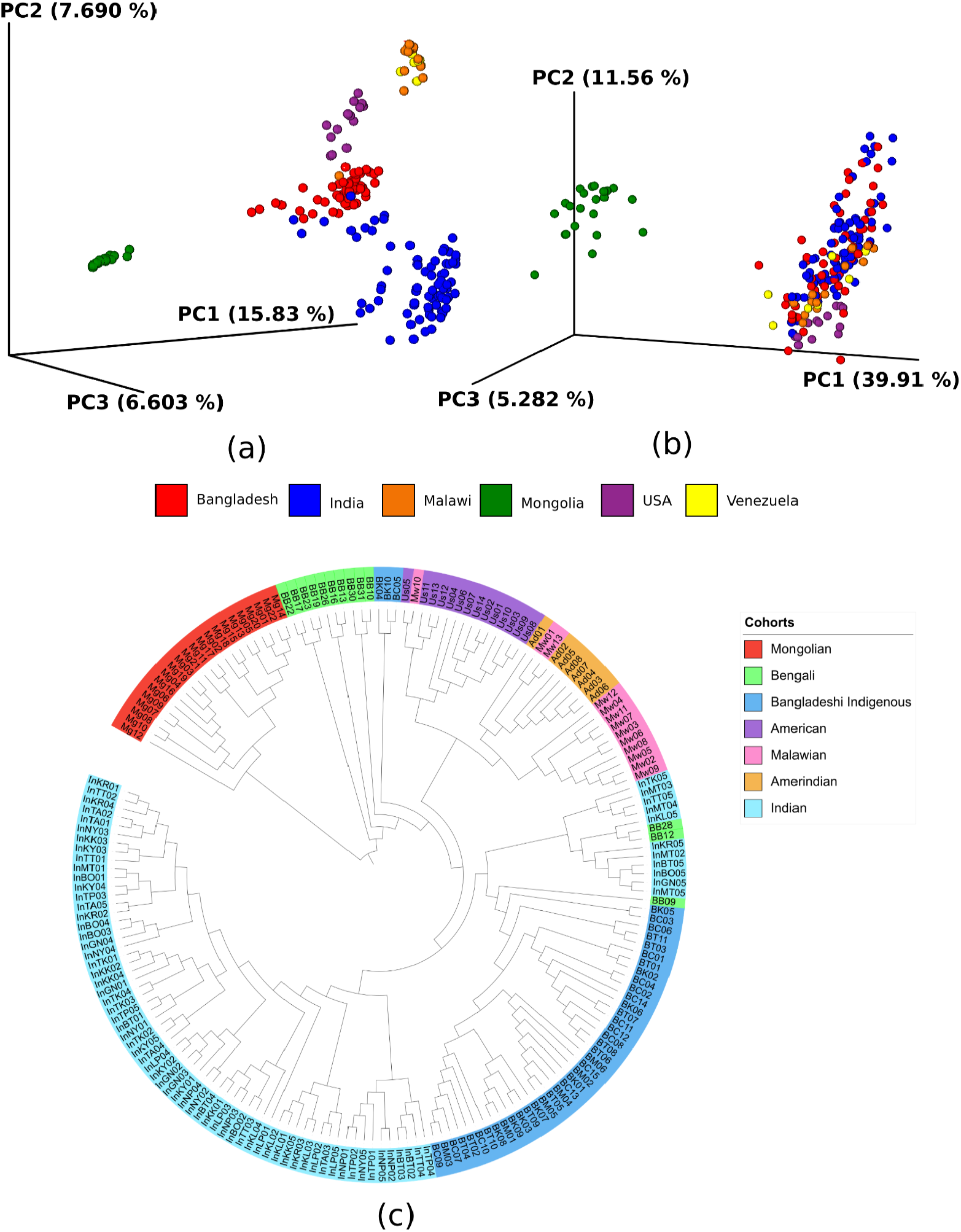
Phylosymbiotic relationships between Bangladeshi communities and other foreign ethnicities. (a) Based on the phylogenetic richness (Unweighted UniFrac distance) of gut microbiota, all Bangladeshi samples clustered in between Mongolian and Indian, Malawian, Amerindian, and American populations. (b) But upon adding phylogenetic quantitative weight (Weighted UniFrac distance), the Bangladeshi population got fully separated from Mongolian samples and clustered with other populations. (c) In terms of gut bacterial structure, the tribal population of Northeast India was more homogeneous with Bangladeshi tribes and clustered together in the UPGMA tree.

### Gut Microbiota Composition of Bangladeshi People is Remarkably Distinct from Other Tropical and Sub-tropical Countries

When compared with the gut microbiome of several other tropical and sub-tropical countries (n=132), the Bangladeshi samples (n=55) formed a vividly distinct cluster (Fig 7). The gut microbes in Bangladeshi samples were not only qualitatively different from others but also phylogenetically different as measured by Faith’s phylogenetic analysis. In terms of beta diversity, Bangladeshis were found to be both qualitatively and quantitively very different from other tropical and sub-tropical countries. This indicates that even though the mainstream Bengali population is different from the indigenous people living in the Chittagong hill tracts located in south-eastern parts of the country, overall the people living within the borders of Bangladesh (collectively referred to as Bangladeshi people) possess a unique gut microbiota composition.

**Fig 07:**
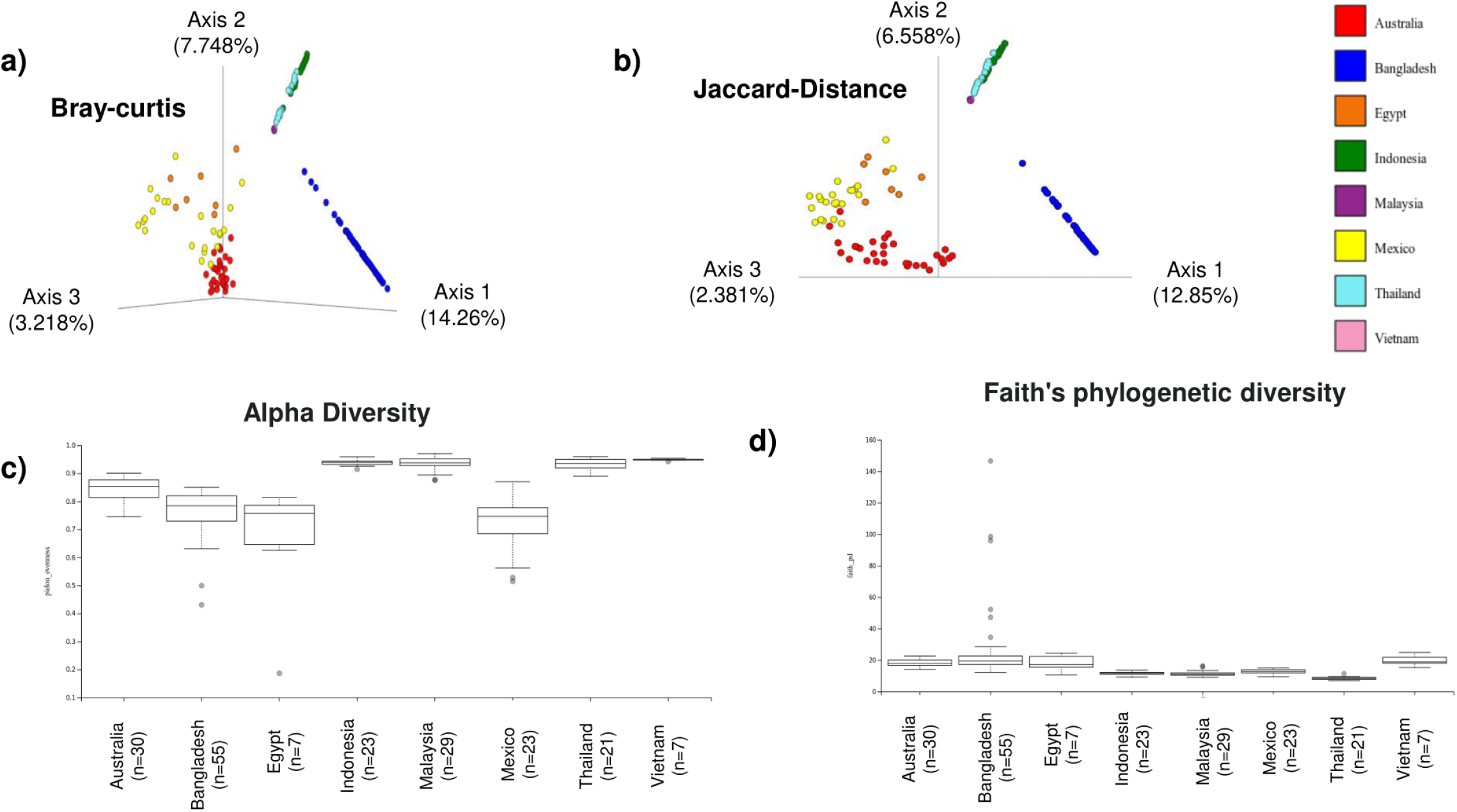
Comparative analysis of the gut microbiome of the Bangladeshi population (n=55) and populations from several tropical and sub-tropical countries (n=132). The parameters used for comparison were (a) Bary-Curtis distance (b) Jaccard Distance (c) Alpha diversity (d) Faith’s phylogenetic diversity

## Discussion

The human gut microbiome is an integral part of human growth and development. The microbiome composition aids in the regulation of the immune system, food digestion, and proper physiology^65^. Ethnicity also plays a vital role in the modulation of the human gut microbiome. Studies on the human microbiome suggest that the typical human gut microbiota is dominated mainly by Bacteroidetes and Firmicutes phyla^66,67^. Their ratio has significant value on human physiology.

According to several studies, the Firmicutes/ Bacteroidetes ratio (F/B) is a crucial biomarker for maintaining normal intestinal physiology^68^. The proportion of Firmicutes and Bacteroidetes can influence our body weight^69,70^. Higher Firmicutes to Bacteroidetes ratio (F/B) is believed to be linked with obesity in humans^71^. In our study, all indigenous cohorts have a median F/B ratio of more than 1.00 and among them, the Tripura tribe had the highest ratio (**Fig. 1d**). In contrast, Bengali populations had a median F/B ratio lower than 1.00, indicating a relatively lower prevalence of Firmicutes phylum in comparison to Bacteroidetes in their gut. The two major representative phyla in the human gut are Bacteroides and Prevotella^72^. People consuming plant-rich diets tend to have a microbiome enriched by the *Prevotella* genus whereas people consuming protein-rich food predominantly have *Bacteroides* in their gut^73^. According to earlier studies, there is a definitive relationship between gut microbiota composition and long-term eating habits^73^. In this study, all cohorts appeared to have an almost similar dispensation of species from the *Prevotella* genus suggesting the frequency of plant-based dietary habits among the people of Bangladesh people (**Fig. 1a**). Contrarily, Chakma people exhibited the highest prevalence of the *Bacteroides* genus compared to other tribes, indicating that the Chakma people consume more animal fat than others and this has shaped their gut microbiome composition. The Marma and Tripura people, on the other hand, still cherish their own traditional agro-based eating habits. Certain intestinal archaea *Methanobrevibacter* and *Methanosphaera* were found in some samples of each cohort (**Fig. 1b**). Previously reported many species from *Methanobrevibacter* and *Methanosphaera* genera had previously been correlated with higher production of methane, causing irritable bowel syndrome (IBS), particularly constipation^74,75^.

The abundance pattern of core features was similar for samples from all indigenous cohorts whereas, in the case of the majority Bengali population, the *Lachnospiraceae, Streptococcaceae Prevotellaceae, Ruminococcaceae*, and *Bacteroidaceae* were the most dominant genus (**Fig. 1c**). For proper interaction between gut microbiota and human health, different species from *Firmicutes* and *Bacteroidetes* phylum play a major role. For instance, our body can not digest resistant fibers. Microbial species from these phyla can break down these carbohydrates through the fermentation process^76^. Many members of *Firmicutes* and *Bacteroidetes* phyla (e.g., *Streptococcaceae, Veillonellaceae, Lachnospiraceae, Ruminococcaceae, Prevotellaceae* families) serve as probiotics and generate short-chain fatty acids (SCFA) like butyrate^77–79^. For example, *Ruminococcus*, a species from the *Ruminococcaceae* family, is one of the three enterotypes of human gut microbiomes, its prevalence in the gut has been connected to starch and dietary fibers containing a long-term diet^80^. A reduction in the abundance of the *Prevotellaceae* population can lead to increased gut permeability and systemic exposure to bacterial endotoxin, which can lead to excessive expression of misfolded synuclein in the colon^81,82^. The decreased abundance of the *Bacteroidaceae* population in the gut has been associated with increased intestinal permeability and thus increases lipopolysaccharide absorption which results in obesity, and inflammation^83,84^. *Enterobacteriaceae* abundance in the gut showed a positive correlation with postural instability and gait difficulty^83^.

The diversity within samples can be evaluated by different alpha diversity metrics, which explain richness, evenness, and also the phylogenetic relationship between microbiota^85^. Having the same alpha diversity indices does not necessarily mean having similar microbial compositions^86^. In this study, the Bengali samples had lower bacterial richness (observed features) than indigenous cohorts, however, they had higher variation within individual samples (**Fig. 2**). As one previous study suggests, ethnic differences might have an influential role in shaping such variation of alpha diversity among cohorts^87^.

On the other hand, beta-diversity metrics estimate between-host microbiome diversity among different metadata categories. In this context, Jaccard distance evaluates the qualitative community dissimilarity, and unweighted UniFrac combines this measure with the phylogenetic relationship. The Bray-Curtis distance, on the other hand, adopts a quantitative approach to appraise dissimilarity between different sample groups, whereas weighted UniFrac includes a phylogenetic correlation in addition to such quantitative metrics^88^. In the current study, it was found that the majority of Bengali samples were distant from Bangladeshi indigenous cohorts in terms of their gut bacterial composition (**Fig. 3a and 3b**). Moreover, among indigenous groups, the Chakma tribe had more distinct microbial features than others (**Fig. 3c**).

Several differentially abundant bacteria were observed in each cohort (**Figure 4**). This differential abundance can play roles in human health and disease^89^. For instance, the Bengali population might face relatively fewer intestinal infections. An increase in the abundance of *Bacteroides caccae*, which was lower in the Bengali population, can promote the erosion of the intestinal mucus layer, which aids in receding inflammation in the intestine. A thinner layer of the intestinal mucus can result in enhanced susceptibility to pathogens^90^. Diarrhea symptoms linked with *Cryptosporidium* infection may be accompanied by a decrease in the abundance of *Megasphaera* species^91^. In the Marma group, the *Lachnospiraceae* family was higher in abundance. The Anaerostipes genus is a member of the *Lachnospiraceae* family and contains gram-positive anaerobic bacteria, which are reported to have a protective role against human colon cancer by producing butyric acid^92^. Moreover, their higher numbers of *Roseburia* species might promote lower glucose intolerance and loss of weight^93^. In the Chakma population, the *Alistipes* genus, which showed higher abundance, is a comparatively new member of the *Bacteroidetes* phylum and it is highly correlated with dysbiosis and disease^94^. In addition, their enhanced *Parabacteroides* genus has been reported to be both beneficial and pathogenic for the human host. *Parabacteroides distasonis*, a gram-negative anaerobic bacteria from this genus, seemed to be linked with antimicrobial resistance as well as pathogenic and probiotic functions in human health^95^. Khyang population had higher levels of *Enterobacteriaceae* that are regarded as commensals in the human gut, along with their beneficial roles, some members of this family, such as *Escherichia, Shigella*, and *Salmonella* spp., can be infectious^96^. Furthermore, their gut microbiome might be less obesogenic due to the abundance of *Oscillospira* species. *Oscillospira* was found to be positively correlated with a low BMI^97^. In the Tripura group, the dominant *Pasteurellaceae* was reported to be more ubiquitous in diabetic retinopathy than in diabetic patients without retinopathy^98^. Their *Haemophilus* predominance is positively associated with depression, cognition, excitement, and negative psychiatric symptoms^99^.

Enrichment of various metabolic pathways was also assessed in this study because the host-microbiome composition has a strong relationship with several biochemical reactions within the host^100^. Bengali samples had a wide variety of pathways functionally enriched by gut microbiome contribution. Among them, DNA replication and repair, translation, amino acid metabolism, glycan biosynthesis, and metabolism, vitamins and cofactor metabolism, nucleotide metabolism, genetic information processing, etc related pathways had higher relative abundance in the Bengali population (**Fig. 5a**). Enrichment of such cellular and genetic processing pathways suggests active cellular activity in Bengali populations. On the other hand, histidine is essential for muscle carnosine synthesis, and its metabolism is associated with physical endurance, recognition memory, neural protection, satiety, and allergic diseases^101,102^. The presence of this metabolic pathway in the Chakma and Marma tribes’ samples indicates that SCFAs play an essential role in the metabolism of their diet (**Fig. 5b and 5c**). L-arginine is suggested to have a protective role against ulcerative colitis and decreased in its biosynthesis can result in several inflammatory bowel diseases^103,104^.

For a better understanding of the underlying mechanism of interaction between humans and the microbiome, an evolutionary perspective can also be helpful^105^. Hence, we assessed the gut microbiome composition of our Bangladeshi cohorts with some geographically related and distant population groups in the context of phylogenetic relationships. The similarity or dissimilarity between different population groups was explored by principal coordinate analysis (PCoA) based on both qualitative and quantitative phylogenetic measures (Unweighted and Weighted UniFrac Distance). In the case of qualitative phylogenetic relationship, Bangladeshi cohorts fall between Mongolian and other populations (North Indian, Amerindian, Malawian, and American) in terms of gut microbiota composition (**Fig. 6a**). But, when the quantitative calculations were added to this relationship, Bangladeshi cohorts moved away from the Mongolian samples and merged with other populations (**Fig. 6b**). Moreover, in the case of UPGMA hierarchical clustering, samples from indigenous people of our country formed clusters close to north Indian tribes (**Fig. 6c**), and most Bengali samples clustered separately. Though the three samples clustered far apart from the other Bengali samples, this can be due to specific health implications of those individuals and should be considered as outliers. Our tribal populations are said to share a common ancestor with the Tibeto-Burman people of Northeast India, and the gut microbial composition-based clustering found in this study supports this conception.

A comparative analysis between Bangladeshi samples as a whole and tropical/sub-tropical samples unveiled the fact that Bangladeshi gut microbiota composition is quite distinct from others (**Fig 7**). Finally, our observation suggests that throughout history, the mix of Tibeto-Burman and Bengali genetics shaped a unique structure of gut-microbiota in indigenous cohorts. Neither tropical populations nor those from Bengal share this type of structure. These groups share similar genetics and indistinguishable gut microbiota amongst themselves but are distinguishable from the mainstream Bengali population. However, urbanization and alteration of food habits will affect the bacterial diversity differently among these populations in the future^106^.

## Conclusion

The study highlights the impact of genetic background and lifestyle on the gut microbiome composition of several ethnicities living in Bangladesh. Although the population of the four indigenous cohorts included in this study resides in the same geographic region of Bangladesh, there are some subtle differences in their gut microbiota profile. But the ethnicity, residence, and lifestyle of Bengali individuals are very different from tribal ones and we found a clear impact of these factors in the microbial community composition. We noticed that among the tribal groups the Chakma tribe had a significant difference in bacterial diversity and contained the most characteristic microbial communities. Moreover, the microbiome content of all cohorts reflected their plant-based diet. It also revealed the animal-fat dominating food habit of Chakma people. Finally, the gut microbiota composition of the aboriginal groups of Bangladesh has also been found to be similar to various genealogically related Indian tribes. Our findings suggest that in addition to ethnic origin, long-term food habits and social circumstances played a prominent role in molding the gut microbiota profile of the Bangladeshi population.

## Acknowledgements

This study was funded under the “Establishment of National Gene Bank Project”, National Institute of Biotechnology (NIB), Ministry of Science and Technology, Government of the People’s Republic of Bangladesh. We would like to thank Md. Amjad Hossain, Ritaren Chakma, Tina Tripura, Barshan Chakma, and Suborna Dash for their assistance with sample and metadata collection.

## Author contributions

I.A., A.R., Z.M.C. and A.B. performed DNA extraction, bioinformatics analyses, and drafted the manuscript. G.D. and S.T. co-ordinated the sampling and metadata collection. C.A.K., K.C.D., M.U.H. edited the manuscript. M.S. supervised the study. All authors have read and approved the manuscript.

## References

1. Ley, R. E., Peterson, D. A. & Gordon, J. I. Ecological and evolutionary forces shaping microbial diversity in the human intestine. Cell 124, 837–848 (2006).

2. Holmes, E., Li, J. V., Marchesi, J. R. & Nicholson, J. K. Gut microbiota composition and activity in relation to host metabolic phenotype and disease risk. Cell Metab. 16, 559–564 (2012).

3. Shreiner, A. B., Kao, J. Y. & Young, V. B. The gut microbiome in health and in disease. Curr. Opin. Gastroenterol. 31, 69–75 (2015).

4. Sekirov, I., Russell, S. L., Antunes, L. C. M. & Finlay, B. B. Gut microbiota in health and disease. Physiol. Rev. 90, 859–904 (2010).

5. Vyas, U. & Ranganathan, N. Probiotics, prebiotics, and synbiotics: gut and beyond. Gastroenterol. Res. Pract. 2012, 872716 (2012).

6. Guarner, F. & Malagelada, J.-R. Gut flora in health and disease. Lancet Lond. Engl. 361, 512–519 (2003).

7. Filippo, C. D. et al. Impact of diet in shaping gut microbiota revealed by a comparative study in children from Europe and rural Africa. Proc. Natl. Acad. Sci. 107, 14691–14696 (2010).

8. Yatsunenko, T. et al. Human gut microbiome viewed across age and geography. Nature 486, 222–227 (2012).

9. Rampelli, S. et al. Metagenome Sequencing of the Hadza Hunter-Gatherer Gut Microbiota. Curr. Biol. CB 25, 1682–1693 (2015).

10. Odamaki, T. et al. Age-related changes in gut microbiota composition from newborn to centenarian: a cross-sectional study. BMC Microbiol. 16, 90 (2016).

11. Hörmannsperger, G., Clavel, T. & Haller, D. Gut matters: microbe-host interactions in allergic diseases. J. Allergy Clin. Immunol. 129, 1452–1459 (2012).

12. Turnbaugh, P. J., Bäckhed, F., Fulton, L. & Gordon, J. I. Diet-induced obesity is linked to marked but reversible alterations in the mouse distal gut microbiome. Cell Host Microbe 3, 213–223 (2008).

13. Dehingia, M. et al. Gut bacterial diversity of the tribes of India and comparison with the worldwide data. Sci. Rep. 5, 18563 (2015).

14. He, Y. et al. Regional variation limits applications of healthy gut microbiome reference ranges and disease models. Nat. Med. 24, 1532–1535 (2018).

15. Schnorr, S. L. et al. Gut microbiome of the Hadza hunter-gatherers. Nat. Commun. 5, 3654 (2014).

16. Clemente, J. C. et al. The microbiome of uncontacted Amerindians. Sci. Adv. 1, e1500183.

17. Gomez, A. et al. Gut Microbiome of Coexisting BaAka Pygmies and Bantu Reflects Gradients of Traditional Subsistence Patterns. Cell Rep. 14, 2142–2153 (2016).

18. Ayeni, F. A. et al. Infant and Adult Gut Microbiome and Metabolome in Rural Bassa and Urban Settlers from Nigeria. Cell Rep. 23, 3056–3067 (2018).

19. Obregon-Tito, A. J. et al. Subsistence strategies in traditional societies distinguish gut microbiomes. Nat. Commun. 6, 6505 (2015).

20. Chakravorty, S., Helb, D., Burday, M., Connell, N. & Alland, D. A detailed analysis of 16S ribosomal RNA gene segments for the diagnosis of pathogenic bacteria. J. Microbiol. Methods 69, 330–339 (2007).

21. Chong, C. W., Pearce, D. A., Convey, P., Yew, W. C. & Tan, I. K. P. Patterns in the distribution of soil bacterial 16S rRNA gene sequences from different regions of Antarctica. Geoderma 181–182, 45–55 (2012).

22. Dethlefsen, L., Huse, S., Sogin, M. L. & Relman, D. A. The Pervasive Effects of an Antibiotic on the Human Gut Microbiota, as Revealed by Deep 16S rRNA Sequencing. PLOS Biol. 6, e280 (2008).

23. Madhavan, A., Sindhu, R., Parameswaran, B., Sukumaran, R. K. & Pandey, A. Metagenome Analysis: a Powerful Tool for Enzyme Bioprospecting. Appl. Biochem. Biotechnol. 183, 636–651 (2017).

24. Sharpton, T. J. An introduction to the analysis of shotgun metagenomic data. Front. Plant Sci. 5, 209 (2014).

25. Quail, M. A. et al. A tale of three next generation sequencing platforms: comparison of Ion Torrent, Pacific Biosciences and Illumina MiSeq sequencers. BMC Genomics 13, 341 (2012).

26. Bangladesh population and housing census, 2011. National report. (Bangladesh Bureau of Statistics, Statistics and Informatics Division, Ministry of Planning, Bangladesh, Government of the People’s Republic of Bangladesh, 2014).

27. Alam, S. M. N. & Naser, M. N. Chapter 2 - Role of traditional foods of Bangladesh in reaching-out of nutrition. in Nutritional and Health Aspects of Food in South Asian Countries (eds. Prakash, J., Waisundara, V. & Prakash, V.) 217–235 (Academic Press, 2020). doi:10.1016/B978-0-12-820011-7.00025-3.

28. Gazi, N. N. et al. Genetic Structure of Tibeto-Burman Populations of Bangladesh: Evaluating the Gene Flow along the Sides of Bay-of-Bengal. PLoS ONE 8, e75064 (2013).

29. Chowdhury, M. S. H. & Miah, Md. D. Housing pattern and food habit of theMro-tribe community in Bangladesh: A forest dependence perspective. J. For. Res. 14, 253–258(2003).

30. Bolyen, E. et al. Reproducible, interactive, scalable and extensible microbiome data science using QIIME 2. Nat. Biotechnol. 37, 852–857 (2019).

31. Wickham, H. et al. Welcome to the Tidyverse. J. Open Source Softw. 4, 1686 (2019).

32. Martin, M. Cutadapt removes adapter sequences from high-throughput sequencing reads. EMBnet.journal 17, 10–12 (2011).

33. Callahan, B. J. et al. DADA2: High-resolution sample inference from Illumina amplicon data. Nat. Methods 13, 581–583 (2016).

34. Pedregosa, F. et al. Scikit-learn: Machine Learning in Python. Mach. Learn. PYTHON 6.

35. McDonald, D. et al. An improved Greengenes taxonomy with explicit ranks for ecological and evolutionary analyses of bacteria and archaea. ISME J. 6, 610–618 (2012).

36. Bokulich, N. A. et al. Optimizing taxonomic classification of marker-gene amplicon sequences with QIIME 2’s q2-feature-classifier plugin. Microbiome 6, 90 (2018).

37. Werner, J. J. et al. Impact of training sets on classification of high-throughput bacterial 16s rRNA gene surveys. ISME J. 6, 94–103 (2012).

38. Bd, O., Nh, B. & Am, P. Interactive metagenomic visualization in a Web browser. BMC Bioinformatics 12, (2011).

39. Hunter, J. D. Matplotlib: A 2D Graphics Environment. Comput. Sci. Eng. 9, 90–95 (2007).

40. Waskom, M. L. seaborn: statistical data visualization. J. Open Source Softw. 6, 3021 (2021).

41. Janssen, S. et al. Phylogenetic Placement of Exact Amplicon Sequences Improves Associations with Clinical Information. mSystems 3, e00021–18.

42. Matsen, F. A., Hoffman, N. G., Gallagher, A. & Stamatakis, A. A Format for Phylogenetic Placements. PLOS ONE 7, e31009 (2012).

43. Weiss, S. et al. Normalization and microbial differential abundance strategies depend upon data characteristics. Microbiome 5, 27 (2017).

44. Halko, N., Martinsson, P.-G., Shkolnisky, Y. & Tygert, M. An Algorithm for the Principal Component Analysis of Large Data Sets. SIAM J. Sci. Comput. 33, 2580–2594 (2011).

45. Vázquez-Baeza, Y., Pirrung, M., Gonzalez, A. & Knight, R. EMPeror: a tool for visualizing high-throughput microbial community data. GigaScience 2, (2013).

46. Kruskal, W. H. & Wallis, W. A. Use of Ranks in One-Criterion Variance Analysis. J. Am. Stat. Assoc. 47, 583–621 (1952).

47. McKinney, W. Data Structures for Statistical Computing in Python. in 56–61 (2010). doi:10.25080/Majora-92bf1922-00a.

48. Bokulich, N. A. et al. q2-longitudinal: Longitudinal and Paired-Sample Analyses of Microbiome Data. mSystems 3, e00219–18 (2018).

49. Anderson, M. J. A new method for non-parametric multivariate analysis of variance. Austral Ecol. 26, 32–46 (2001).

50. Mandal, S. et al. Analysis of composition of microbiomes: a novel method for studying microbial composition. Microb. Ecol. Health Dis. 26, 27663 (2015).

51. Lin, H. & Peddada, S. D. Analysis of microbial compositions: a review of normalization and differential abundance analysis. Npj Biofilms Microbiomes 6, 1–13 (2020).

52. Segata, N. et al. Metagenomic biomarker discovery and explanation. Genome Biol. 12, R60 (2011).

53. McNally, C. P., Eng, A., Noecker, C., Gagne-Maynard, W. C. & Borenstein, E. BURRITO: An Interactive Multi-Omic Tool for Visualizing Taxa–Function Relationships in Microbiome Data. Front. Microbiol. 9, 365 (2018).

54. Langille, M. G. I. et al. Predictive functional profiling of microbial communities using 16S rRNA marker gene sequences. Nat. Biotechnol. 31, 814–821 (2013).

55. Kanehisa, M., Sato, Y., Kawashima, M., Furumichi, M. & Tanabe, M. KEGG as a reference resource for gene and protein annotation. Nucleic Acids Res. 44, D457–D462 (2016).

56. Rognes, T., Flouri, T., Nichols, B., Quince, C. & Mahé, F. VSEARCH: a versatile open source tool for metagenomics. PeerJ 4, e2584 (2016).

57. DeSantis, T. Z. et al. Greengenes, a Chimera-Checked 16S rRNA Gene Database and Workbench Compatible with ARB. Appl. Environ. Microbiol. 72, 5069–5072 (2006).

58. Meyer, F. et al. The metagenomics RAST server – a public resource for the automatic phylogenetic and functional analysis of metagenomes. BMC Bioinformatics 9, 386 (2008).

59. Zhang, J. et al. Mongolians core gut microbiota and its correlation with seasonal dietary changes. Sci. Rep. 4, 5001 (2014).

60. Amir, A. et al. Deblur Rapidly Resolves Single-Nucleotide Community Sequence Patterns. mSystems 2, e00191-16 (2017).

61. Bokulich, N. A. et al. Quality-filtering vastly improves diversity estimates from Illumina amplicon sequencing. Nat. Methods 10, 57–59 (2013).

62. Mantel, N. The Detection of Disease Clustering and a Generalized Regression Approach. Cancer Res. 27, 209–220 (1967).

63. Michener, C. D. & Sokal, R. R. A Quantitative Approach to a Problem in Classification. Evolution 11, 130–162 (1957).

64. Letunic, I. & Bork, P. Interactive Tree Of Life (iTOL) v5: an online tool for phylogenetic tree display and annotation. Nucleic Acids Res. 49, W293–W296 (2021).

65. Guinane, C. M. & Cotter, P. D. Role of the gut microbiota in health and chronic gastrointestinal disease: understanding a hidden metabolic organ. Ther. Adv. Gastroenterol. 6, 295–308 (2013).

66. Ley, R. E. Prevotella in the gut: choose carefully. Nat. Rev. Gastroenterol. Hepatol. 13, 69–70 (2016).

67. Lozupone, C. A., Stombaugh, J. I., Gordon, J. I., Jansson, J. K. & Knight, R. Diversity, stability and resilience of the human gut microbiota. Nature 489, 220–230 (2012).

68. Stojanov, S., Berlec, A. & Štrukelj, B. The Influence of Probiotics on the Firmicutes/Bacteroidetes Ratio in the Treatment of Obesity and Inflammatory Bowel disease. Microorganisms 8, 1715 (2020).

69. Crovesy, L., Masterson, D. & Rosado, E. L. Profile of the gut microbiota of adults with obesity: a systematic review. Eur. J. Clin. Nutr. 74, 1251–1262 (2020).

70. Tseng, C.-H. & Wu, C.-Y. The gut microbiome in obesity. J. Formos. Med. Assoc. 118, S3–S9 (2019).

71. Ley, R. E., Turnbaugh, P. J., Klein, S. & Gordon, J. I. Human gut microbes associated with obesity. Nature 444, 1022–1023 (2006).

72. Thursby, E. & Juge, N. Introduction to the human gut microbiota. Biochem. J. 474, 1823–1836 (2017).

73. Wu, G. D. et al. Linking Long-Term Dietary Patterns with Gut Microbial Enterotypes. Science 334, 105–108 (2011).

74. Ghoshal, U., Shukla, R., Srivastava, D. & Ghoshal, U. C. Irritable Bowel Syndrome, Particularly the Constipation-Predominant Form, Involves an Increase in Methanobrevibacter smithii, Which Is Associated with Higher Methane Production. Gut Liver 10, 932–938 (2016).

75. Guindo, C. O., Davoust, B., Drancourt, M. & Grine, G. Diversity of Methanogens in Animals’ Gut. Microorganisms 9, 13 (2020).

76. Ramakrishna, B. S. Role of the gut microbiota in human nutrition and metabolism. J. Gastroenterol. Hepatol. 28, 9–17 (2013).

77. Venkat, H. K., Sahu, N. P. & Jain, K. K. Effect of feeding Lactobacillus-based probiotics on the gut microflora, growth and survival of postlarvae of Macrobrachium rosenbergii (de Man). Aquac. Res. 35, 501–507 (2004).

78. Parada Venegas, D. et al. Short Chain Fatty Acids (SCFAs)-Mediated Gut Epithelial and Immune Regulation and Its Relevance for Inflammatory Bowel Diseases. Front. Immunol. 10, 277 (2019).

79. Qin, J. et al. A human gut microbial gene catalogue established by metagenomic sequencing. Nature 464, 59–65 (2010).

80. Arumugam, M. et al. Enterotypes of the human gut microbiome. Nature 473, 174–180 (2011).

81. Forsythe, P., Bienenstock, J. & Kunze, W. A. Vagal Pathways for Microbiome-Brain-Gut Axis Communication. in Microbial Endocrinology: The Microbiota-Gut-Brain Axis in Health and Disease (eds. Lyte, M. & Cryan, J. F.) 115–133 (Springer, 2014). doi:10.1007/978-1-4939-0897-4_5.

82. Niehaus, I. & Lange, J. H. Endotoxin: is it an environmental factor in the cause of Parkinson’s disease? Occup. Environ. Med. 60, 378 (2003).

83. Chakraborti, C. K. New-found link between microbiota and obesity. World J. Gastrointest. Pathophysiol. 6, 110–119 (2015).

84. Aguilar, T. et al. Gut Bacterial Families Are Associated with Body Composition and Metabolic Risk Markers in School-Aged Children in Rural Mexico. Child. Obes. 16, 358–366 (2020).

85. Liu, Y.-X. et al. A practical guide to amplicon and metagenomic analysis of microbiome data. Protein Cell 12, 315–330 (2021).

86. Walters, K. E. & Martiny, J. B. H. Alpha-, beta-, and gamma-diversity of bacteria varies across habitats. PLOS ONE 15, e0233872 (2020).

87. Liu, K. et al. Ethnic Differences Shape the Alpha but Not Beta Diversity of Gut Microbiota from School Children in the Absence of Environmental Differences. Microorganisms 8, 254 (2020).

88. Lozupone, C. A., Hamady, M., Kelley, S. T. & Knight, R. Quantitative and Qualitative β Diversity Measures Lead to Different Insights into Factors That Structure Microbial Communities. Appl. Environ. Microbiol. 73, 1576–1585 (2007).

89. Gilbert, J. A. et al. Current understanding of the human microbiome. Nat. Med. 24, 392–400 (2018).

90. Desai, M. S. et al. A Dietary Fiber-Deprived Gut Microbiota Degrades the Colonic Mucus Barrier and Enhances Pathogen Susceptibility. Cell 167, 1339–1353.e21 (2016).

91. Carey, M. A. et al. Megasphaera in the Stool Microbiota Is Negatively Associated With Diarrheal Cryptosporidiosis. Clin. Infect. Dis. 73, e1242–e1251 (2021).

92. Ai, D. et al. Identifying Gut Microbiota Associated With Colorectal Cancer Using a Zero-Inflated Lognormal Model. Front. Microbiol. 10, 826 (2019).

93. Falalyeyeva, T. et al. 2.18 - Gut Microbiota Interactions With Obesity. in Comprehensive Gut Microbiota (ed. Glibetic, M.) 201–219 (Elsevier, 2022). doi:10.1016/B978-0-12-819265-8.00030-9.

94. Parker, B. J., Wearsch, P. A., Veloo, A. C. M. & Rodriguez-Palacios, A. The Genus Alistipes: Gut Bacteria With Emerging Implications to Inflammation, Cancer, and Mental Health. Front. Immunol. 11, 906 (2020).

95. Ezeji, J. C. et al. Parabacteroides distasonis: intriguing aerotolerant gut anaerobe with emerging antimicrobial resistance and pathogenic and probiotic roles in human health. Gut Microbes 13, 1922241 (2021).

96. Kang, E. et al. Enterobacteria and host resistance to infection. Mamm. Genome 29, 558–576 (2018).

97. Chen, Y. et al. High Oscillospira abundance indicates constipation and low BMI in the Guangdong Gut Microbiome Project. Sci. Rep. 10, 9364 (2020).

98. Huang, Y. et al. Dysbiosis and Implication of the Gut Microbiota in Diabetic Retinopathy. Front. Cell. Infect. Microbiol. 11, (2021).

99. Zhu, C. et al. Association Between Abundance of Haemophilus in the Gut Microbiota and Negative Symptoms of Schizophrenia. Front. Psychiatry 12, (2021).

100. Yadav, M., Verma, M. K. & Chauhan, N. S. A review of metabolic potential of human gut microbiome in human nutrition. Arch. Microbiol. 200, 203–217 (2018).

101. Artioli, G. G., Gualano, B., Smith, A., Stout, J. & Lancha, A. H. Role of beta-alanine supplementation on muscle carnosine and exercise performance. Med. Sci. Sports Exerc. 42, 1162–1173 (2010).

102. Acuña, I. et al. Rapid and simultaneous determination of histidine metabolism intermediates in human and mouse microbiota and biomatrices. BioFactors 48, 315–328 (2022).

103. Ren, W. et al. Serum Amino Acids Profile and the Beneficial Effects of L-Arginine or L-Glutamine Supplementation in Dextran Sulfate Sodium Colitis. PLOS ONE 9, e88335 (2014).

104. Vich Vila, A. et al. Gut microbiota composition and functional changes in inflammatory bowel disease and irritable bowel syndrome. Sci. Transl. Med. 10, eaap8914 (2018).

105. Davenport, E. R. et al. The human microbiome in evolution. BMC Biol. 15, 127 (2017).

106. Teyssier, A. et al. Diet contributes to urban-induced alterations in gut microbiota: experimental evidence from a wild passerine. Proc. R. Soc. B Biol. Sci. 287, 20192182 (2020).

107. Penington, J. S. et al. Influence of fecal collection conditions and 16S rRNA gene sequencing at two centers on human gut microbiota analysis. Sci. Rep. 8, 4386 (2018).

108. Radwan, S. et al. A comparative study of the gut microbiome in Egyptian patients with Type I and Type II diabetes. PLOS ONE 15, e0238764 (2020).

109. Therdtatha, P. et al. Gut Microbiome of Indonesian Adults Associated with Obesity and Type 2 Diabetes: A Cross-Sectional Study in an Asian City, Yogyakarta. Microorganisms 9, 897 (2021).

110. Dwiyanto, J. et al. Ethnicity influences the gut microbiota of individuals sharing a geographical location: a cross-sectional study from a middle-income country. Sci. Rep. 11, 2618 (2021).

111. Kitten, A. K., Ryan, L., Lee, G. C., Flores, B. E. & Reveles, K. R. Gut microbiome differences among Mexican Americans with and without type 2 diabetes mellitus. PLoS ONE 16, e0251245 (2021).

112. La-ongkham, O., Nakphaichit, M., Nakayama, J., Keawsompong, S. & Nitisinprasert, S. Age-related changes in the gut microbiota and the core gut microbiome of healthy Thai humans. 3 Biotech 10, 276 (2020).

113. Hoang, H. T. et al. Metagenomic 16S rDNA amplicon data of microbial diversity of guts in Vietnamese humans with type 2 diabetes and nondiabetic adults. Data Brief 34, 106690(2020).

